# In turkeys, unlike chickens, the non-structural NS1 protein does not play a significant role in the replication and tissue tropism of the H7N1 avian influenza virus

**DOI:** 10.1101/2024.01.29.577768

**Authors:** Maryna Kuryshko, Maria Landmann, Christine Luttermann, Reiner Ulrich, Elsayed M. Abdelwhab

**Affiliations:** Institute of Molecular Virology and Cell Biology, Friedrich-Loeffler-Institut, Federal Research Institute for Animal Health, Greifswald-Insel Riems, Germany; Institute of Veterinary Pathology, Faculty of Veterinary Medicine, Leipzig University, Leipzig, Germany; Institute of Immunology, Friedrich-Loeffler-Institut, Federal Research Institute for Animal Health, Greifswald-Insel Riems, Germany

**Keywords:** Virulence determinants, avian influenza virus, turkeys, NS1, interferon, replication, tissue tropism

## Abstract

The economic losses caused by high pathogenicity (HP) avian influenza viruses (AIV) in poultry industry worldwide are enormous. Although chickens and turkeys are two closely related Galliformes, turkeys are thought to be a bridging host for the adaptation of AIV from wild birds to poultry because of their high susceptibility to AIV infections. HPAIV evolve from low pathogenicity (LP) AIV after circulation in poultry through mutations in different viral proteins, including the non-structural protein (NS1), a major interferon (IFN) antagonist of AIV. At present, it is largely unknown whether the virulence determinants of HPAIV are the same in turkeys and chickens. Previously, we showed that mutations in the NS1 of HPAIV H7N1 significantly reduced viral replication in chickens in vitro and in vivo. Here, we investigated the effect of NS1 on the replication and virulence of HPAIV H7N1 in turkeys after inoculation with recombinant H7N1 carrying a naturally truncated wild-type NS1 (with 224 amino-acid “aa” in length) or an extended NS1 with 230-aa similar to the LP H7N1 ancestor. There were no significant differences in multiple-cycle viral replication or in the efficiency of NS1 to block IFN induction in cell culture. Similarly, all viruses were highly virulent in turkeys and replicated at similar levels in various organs and swabs collected from inoculated turkeys. These results suggest that NS1 does not play a role in the virulence or replication of HPAIV H7N1 in turkeys and further indicate that the genetic determinants of HPAIV differ in these two closely related galliform species.

## Introduction

Avian influenza viruses (AIV) belong to the family *Orthomyxoviridae* and the genus Influenza A virus (IAV). AIV are further subdivided into 16 HA and 9 NA subtypes according to variations in the surface glycoprotein, haemagglutinin (HA or H) and neuraminidase (NA or N). AIV infect a wide range of bird species. Wild birds are the natural reservoir for AIV. Infection of wild birds with AIV is, with rare exceptions, asymptomatic, while in poultry it can cause high mortality with huge economic impact. AIV usually has two types of pathogenicity: low pathogenicity (LP) and high pathogenicity (HP) forms ^1^. HPAIV H5 and H7 subtypes evolve from LP ancestors by acquisition of point mutations in different proteins or by reassortment (swapping) of gene segments between two different AIVs infecting the same cell ^1^. Although the acquisition of a polybasic cleavage site in HA is a major determinant of AIV pathogenicity ^2^, other genes contribute to the virulence and pathogenesis of HPAIV ^3^. Compared to chickens, very little is known about the genetic determinants for replication and virulence of HPAIV in turkeys, the second most kept poultry species in Europe and USA after chickens.

Non-structural protein 1 (NS1) has been described to contribute to virulence in chickens and ducks ^4, 5^. NS1 is encoded by the shortest segment of the AIV genome with a typical length of 230 amino acids (aa). NS1 is a multifunctional protein consisting of an RNA binding domain (RBD) linked by a short linker (LR) to an effector domain (ED) ^6^. The RBD of the NS1 protein binds to different types of RNA, preventing the activation of host cell sensors for foreign viral RNAs. The ED is responsible for interacting with various host factors via multiple motifs to trigger different pathways of the innate immune response following AIV infection ^7, 8^. Variations in stop codons in the ED, resulting in truncation of the carboxyl terminus (CTE), may lead to variations in the size of NS1. Truncation of CTE of NS1 has been described to affect its main function, interferon antagonism ^7, 8^. In 1999, H7N1 LPAIV emerged in March and HPAIV in December in Italy. Several genetic changes accompanied the shift to HPAIV and have been described previously ^9^. Truncation of 6-aa in the NS1 CTE and three mutations in the ED accompanied the evolution of the Italian HPAIV H7N1. Previously, we showed that the extension of the NS1 of the Italian HPAIV H7N1 (from 224-aa to 230-aa, similar to the LP precursor) reduced HPAIV replication and cell/tissue tropism in chickens ^10^. Likewise, reduced virulence was observed in chicken embryos ^11^. Although turkeys were the most affected species among the 13 million dead birds during the Italian outbreak ^12^, little is known about the effect of NS1 CTE truncation on AIV virulence in turkeys. Here, we investigated the effect of NS1 length variation on virus replication *in vitro* and virulence and transmission in experimentally inoculated turkeys.

## Results

### Sequence analysis suggested that turkeys, unlike chickens, are infected with H7N1 AIV carrying variable NS1 with different lengths

We analyzed the NS1 sequences of Italian H7N1 available in the GISAID on 16-01-2024 (n= 124) isolated from turkeys (n= 71), chickens (n= 35) and other birds (n= 18). As previously reported ^9, 10^, all H7N1 LPAIV in 1999 contained the full-length NS1 with 230-aa. Interestingly, we found that almost all non-turkey isolates in 2000 contained a short NS1 with 224-aa in length. Conversely, turkey isolates in 2000 (n= 27) showed variations in the NS1 length and possessed NS1 with 230-aa (n= 1/27, an LP), 224-aa (n= 14/27), 220-aa (n=11/27) or 202-aa (n=1/27) (Fig. 1A). *These results suggest that, in contrast to chickens, there may be no positive selection or advantage for NS1 CTE variations occurred in H7N1 isolated from turkeys*.

**Fig. 1:**
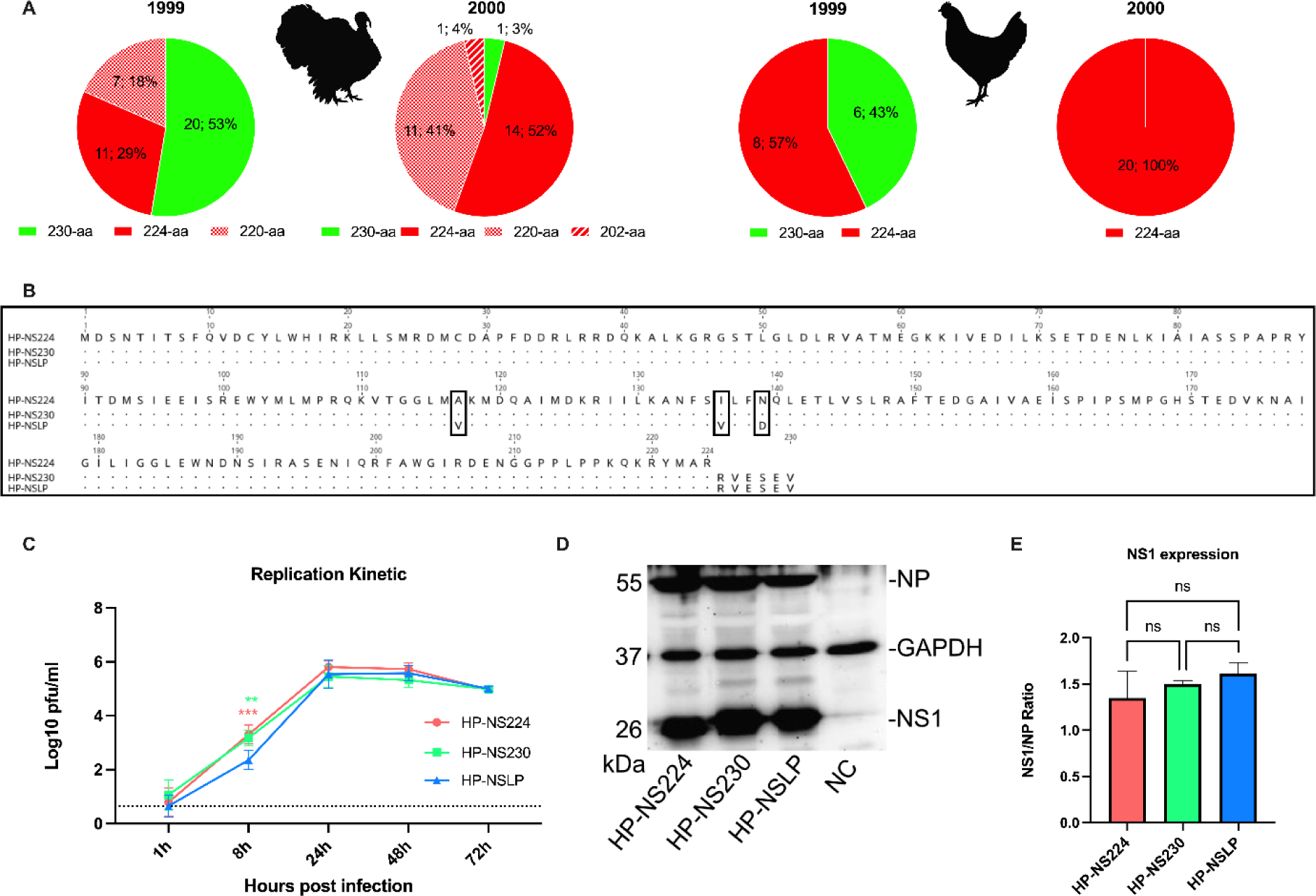
Sequence analysis and in vitro characterization of recombinant H7N1 viruses. (A) Sequence analysis of NS1 of H7N1 from turkeys (n= 65) and chickens (n= 28) in 1999-2000 deposited in GISAID (retrieval date 16-01-2024). Variations in NS1 length are shown as number and percentage of total sequences analyzed in different years. Sequence analysis was performed using Geneious Prime. (B) Alignment and amino acid differences of the NS1 protein of the viruses used for this study. (C) Turkey embryonic kidney (TEK) cells were infected at MOI 0.001 and virus titers were determined at 1, 8, 24, 48 and 72 h post infection (hpi) by plaque assay performed on MDCK II cells. Titers were calculated as pfu/ml and are presented as the mean and standard deviation of three independent experiments performed in duplicate. Data were analyzed by one-way ANOVA with post hoc Tukey test. Asterisks indicate significant differences (*= p< 0.05, **= p < 0.01, ***= p < 0.001, ****= p< 0.0001); ns= no significant differences. (D) NS1 protein was detected after infection of TEK cells at a MOI of 0.1 for 24 h at 37°C. Detection was performed using rabbit polyclonal sera (D. Marc, INRAE, Nouzilly, France) and ECL substrate. The Western blot image was acquired using Quantity One software version 4.4 (Biorad, Germany). (E) Image J was used to calculate NS1 expression levels and data are presented as NS1/NP ratio.

### Recombinant viruses and mutants

To assess the effect of NS1 length variation on the fitness of the Italian HPAIV H7N1, we used 3 recombinant viruses, generated in a previous study ^10^. Each of these viruses contains 7 gene segments (segments 1 to 7) of the Italian H7N1 HPAIV and variable NS segments (Fig. 1B): HP-NS224 (contains NS from HP with NS1 of 224-aa length), HP-NS230 (NS from HP with extended CTE similar to LP NS1) and HP-NSLP (contains NS from LP virus with 230aa length and 3-aa substitutions in the ED: V117A, V136I and D139I). Prior to use, all viruses were sequenced and found to have no undesirable mutations.

### NS1 did not significantly affect multi-cycle replication of HPAIV H7N1 in primary turkey cells

A previous study showed that NS1 CTE and ED affected the replication of an Italian H7N1 virus in chicken cells ^11^, while another study did not find any difference in the replication of LP H7N1 with similar NS1-230 or NS1-224 in chicken or duck cells ^13^. Here, we investigated the influence of NS1 on HPAIV H7N1 replication in freshly prepared primary turkey embryo kidney (TEK) cells infected at a multiplicity of infection (MOI) of 0.001 for single cycle (8 hours post infection “hpi”) and multiple cycle replication (24, 48 and 72 hpi) (Fig. 1C). Virus titer was determined by plaque assay on MDCKII cells. All viruses reached their maximum replication rate at 24 hpi. There were no significant differences in virus titers 24, 48 or 72 hpi. At 8 hpi, HP-NSLP had a significantly lower titer compared to HP-NS224 and HP-NS230 (p < 0.01). *These results indicate that NS1 did not significantly affect the multiple-cycle replication of HPAIV H7N1 in primary turkey cells*.

### Variations in the NS1 protein did not significantly affect NS1 expression levels in primary turkey cells

To assess the effect of changes in CTE or ED on the expression of NS1 in turkey cells, TEK cells were infected with HP-NS224, HP-NS230 or HP-NSLP at a MOI of 0.1 pfu for 24 h (Fig. 1D-E). Cell lysates were prepared and protein expression was assessed by Western blot. GAPDH protein and H7N1 nucleoprotein (NP) were used as internal controls. As expected, the molecular mass of NS1 correlated with the length of the NS1 protein. However, *there were no significant differences in the expression levels of the NS1 variants in TEK cells under these experimental settings*.

### All NS1 proteins were comparably efficient at blocking IFN-α and IFN-β mRNA induction in cell culture

Previous reports indicated that HPAIV NS1 was more efficient at inhibiting IFN-induction of similar Italian HPAIV H7N1 in chicken fibroblast cells [<u>16</u>]. To investigate whether the length of NS1 had an effect on the IFN antagonism of our H7N1 viruses, we used a Firefly-luciferase reporter assay after transfection of human (HEK293T) and avian (DF1) cells (Fig. 2). Attempts to transfect primary TEK cells were unsuccessful. Data were normalized to signals induced in cells transfected with empty vectors. *All three NS1 pCAGGS expression plasmids were comparably efficient in blocking IFN-α and IFN-β mRNA induction in cells triggered by various avian and human factors like MDA5, Trif or IRF7*.

**Fig. 2:**
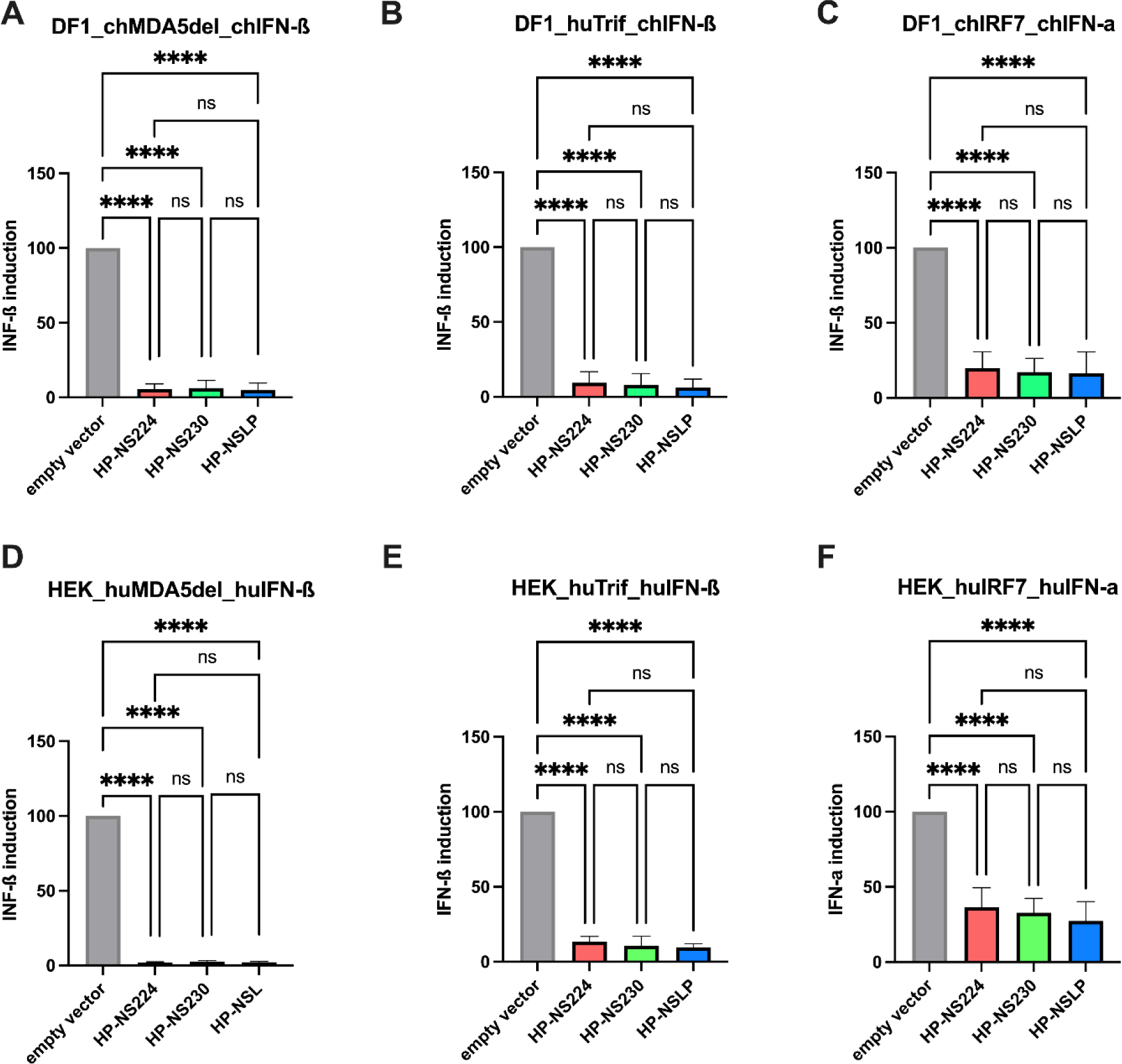
Inhibition of type I interferon induction in avian and human cell lines after transfection with H7N1 NS1. The efficiency of NS1 to block IFN-a and -b mRNA induction was investigated in avian (DF1) (A-C) and human (HEK293T) (D-F) cells using a double reporter luciferase assay. Results are expressed as fold change of IFN-I promotor induction relative to the signal of the indicated trigger for the empty vector control. Asterisks indicate significant differences (*= p< 0.05, **= p < 0.01, ***= p < 0.001, ****= p< 0.0001). ch, chicken; hu, human; ns= no significant differences.

### Variations in NS1 did not significantly affect the high virulence or transmissibility of H7N1 AIV in turkeys

Previous studies have shown that NS1 mutations reduce the virulence of an H5N1 HPAIV in chickens ^5, 14^, while others have found no variations in the high virulence of H7N1 in chickens ^10^. Furthermore, no data are available on the influence of NS1 on the transmission of HPAIV in turkeys. Therefore, we oculonasally inoculated 6-week-old turkeys (n=10) with recombinant HPAIV H7N1 and 1-day post inoculation (dpi) naïve turkeys (n=5) were added to each group to assess turkey-to-turkey transmission. Morbidity, mortality and clinical scores (ranging from 0 = avirulent to 3 = highly virulent) were recorded daily for 10 days (Table 1; Fig. 3A, B). Almost all birds developed clinical signs such as ruffled feathers, mild depression and neurological disorders starting 2 dpi and progressed moderately 3 dpi. All HP-NSLP-inoculated birds had died 4 dpi and all HP-NS230 and HP-NS224-inoculated groups had died 5 dpi. The contact birds in all three groups started to show clinical signs 3 dpi and were dead 5 dpi in the HP-NSLP and HP-NS230 groups and 6 dpi in the HP-NS224 group. No significant difference in pathogenicity index (PI) was observed for all viruses, with mean clinical scores of 2.44, 2.42 and 2.48, respectively (Table 1; Fig. 3A, B). *Taken together, NS1 did not significantly affect the high virulence or efficient transmission of HPAIV H7N1 in turkeys*.

**Fig. 3:**
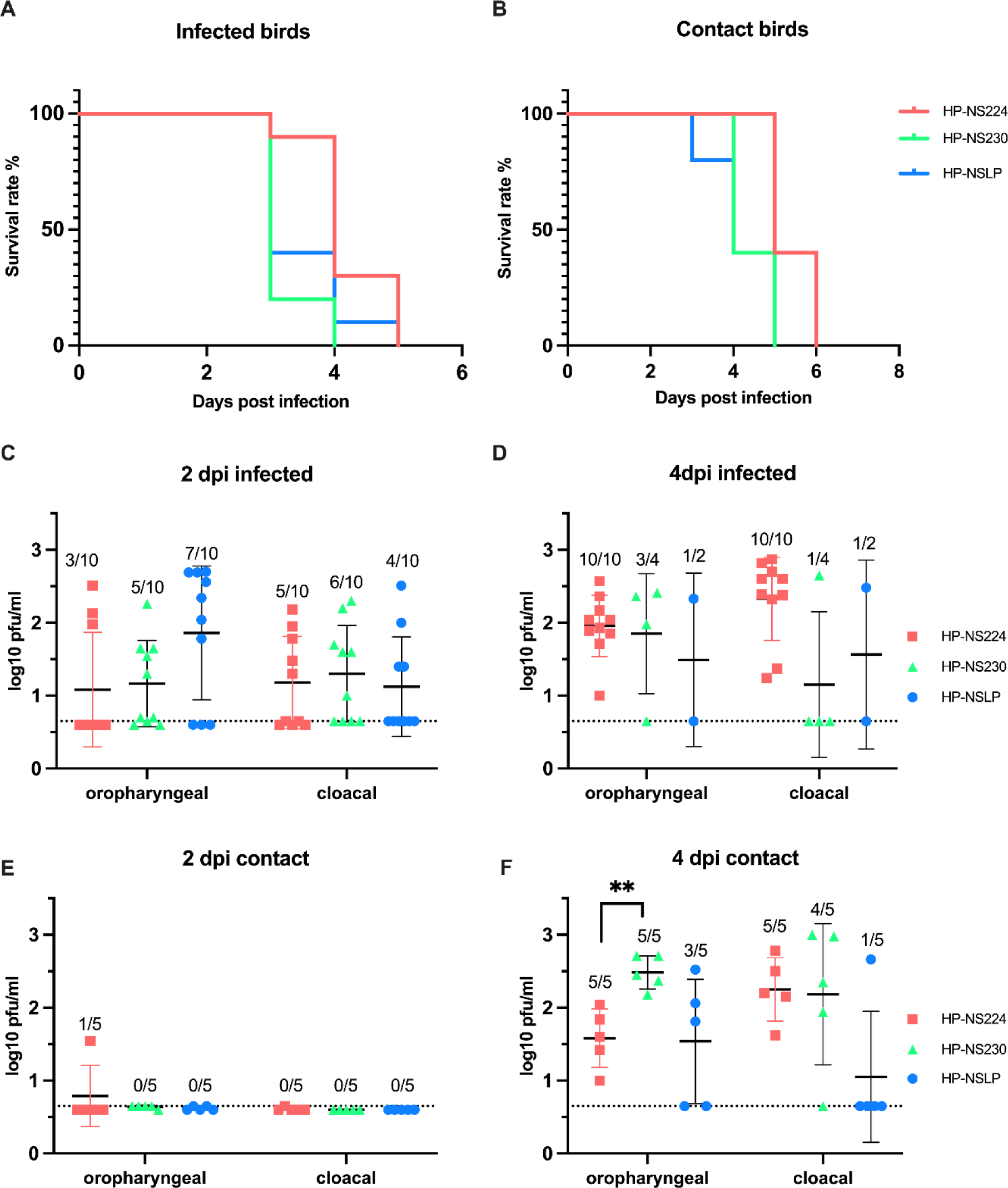
Effect of NS1 gene segment on survival and virus shedding after infection of turkeys with recombinant H7N1 viruses. The survival rate is shown for both inoculated- (A) and contact-turkeys (B). Detection of viral load in swab samples from infected (C, D) and contact (E, F) birds. Oropharyngeal and cloacal swabs were collected 2- and 4-days post infection (dpi) and viral titers were measured by plaque assay using MDCKII cells. Asterisks indicate significant difference, **= p < 0.01. Dashed lines (C, D, E, F) indicate the predicted limit of detection of the plaque assay.

**Table 1.** Virulence of recombinant H7N1 viruses in inoculated and contact turkeys.

| Viruses | Inoculated turkeys |  |  | Contact turkeys |  |
| --- | --- | --- | --- | --- | --- |
|  | Mortality | PI | MTD | Mortality | MTD |
| HP-NS224 | 10/10 | 2.44 | 4.2 | 5/5 | 5.4 |
| HP-NS230 | 10/10 | 2.42 | 3.2 | 5/5 | 4.4 |
| HP-NSLP | 10/10 | 2.48 | 3.5 | 5/5 | 4.2 |

Ten turkeys were inoculated with $10^{4.5}$ pfu/ml of the indicated viruses. To assess bird-to-bird transmission, 5 naive birds were added to each inoculated group at 1-day post infection (dpi). Mortality refers to dead animals/total inoculated. Pathogenicity index (PI) from 0 (avirulent) to 3 (highly virulent) and mean time to death (MTD) per day after infection was calculated as the sum of daily arithmetic means divided by 10, the number of days observed. MTD in contact turkeys was calculated from the dpi of the primary inoculated birds.

### NS1 did not affect viral shedding in primary infected turkeys, but extension of NS1 CTE increased viral shedding in co-housed turkeys

Our previous experiment in chickens ^10^ clearly showed that HP-NS1 extension reduced HPAIV H7N1 shedding and suggested a role for NS1 in virus transmission in chickens. We therefore measured infectious virus in cloacal and oropharyngeal swabs 2 and 4 dpi. The inoculated turkeys shed comparable amount of virus at 2 and 4 dpi with no statistical differences between groups. In the sentinel turkeys, virus titers were comparable to those of the primary inoculated birds 4 dpi. The only significant difference was that HP-NS230 was shed in the oropharyngeal swabs at significantly higher levels (∼10 times higher) than HP-NS224 at 4 dpi (Fig. 3C-F). *Collectively, these results suggest that NS1 did not significantly affect virus replication in primarily inoculated turkeys or transmission to co-housed turkeys*.

### NS1 did not greatly affect the histopathological lesions or tropism in inoculated turkeys

To assess the distribution of H7N1 matrix 1 (M1) antigen as an indicator for virus replication, organ samples collected 4 dpi were subjected to immunohistochemical examination (Fig. 4) similar to our previous chicken experiment ^10^. No tropism to blood vessel endothelium was observed in any of the turkeys, except for a few focal signals, mostly in the nasal cavity of HP-NS224 and HP-NSLP inoculated turkeys (Fig. 4A). Conversely, semiquantitative assessment of M1 protein showed for all three viruses comparable, often multifocal or diffuse distribution of antigen in the parenchyma of different organs such as brain, heart, kidney, pancreas and trachea, but no or only rarely focal parenchymal antigen in the duodenum, proventriculus, liver, lung or skin (Fig. 4B, Fig. 5). The distribution of M1 antigen in cardiomyocytes was more widespread in HP-NS230 than in the other two groups (Fig. 4B). For all three viruses, similar levels of necrosis were also observed mainly in the brain, heart, kidney, nasal cavity, pancreas and trachea, ranging from mild to severe (Fig. 4C). Depletion was observed in lymphoid organs in all turkeys examined, particularly in those inoculated with HP-NSLP, followed by HP-NS230 and HP-NS224 (Fig. 4D). *These results indicate that variations in NS1 did not significantly affect virus distribution or histopathological changes in the major organs (e.g. brain, heart, kidney, spleen), which is consistent with clinical examination*.

**Fig. 4:**
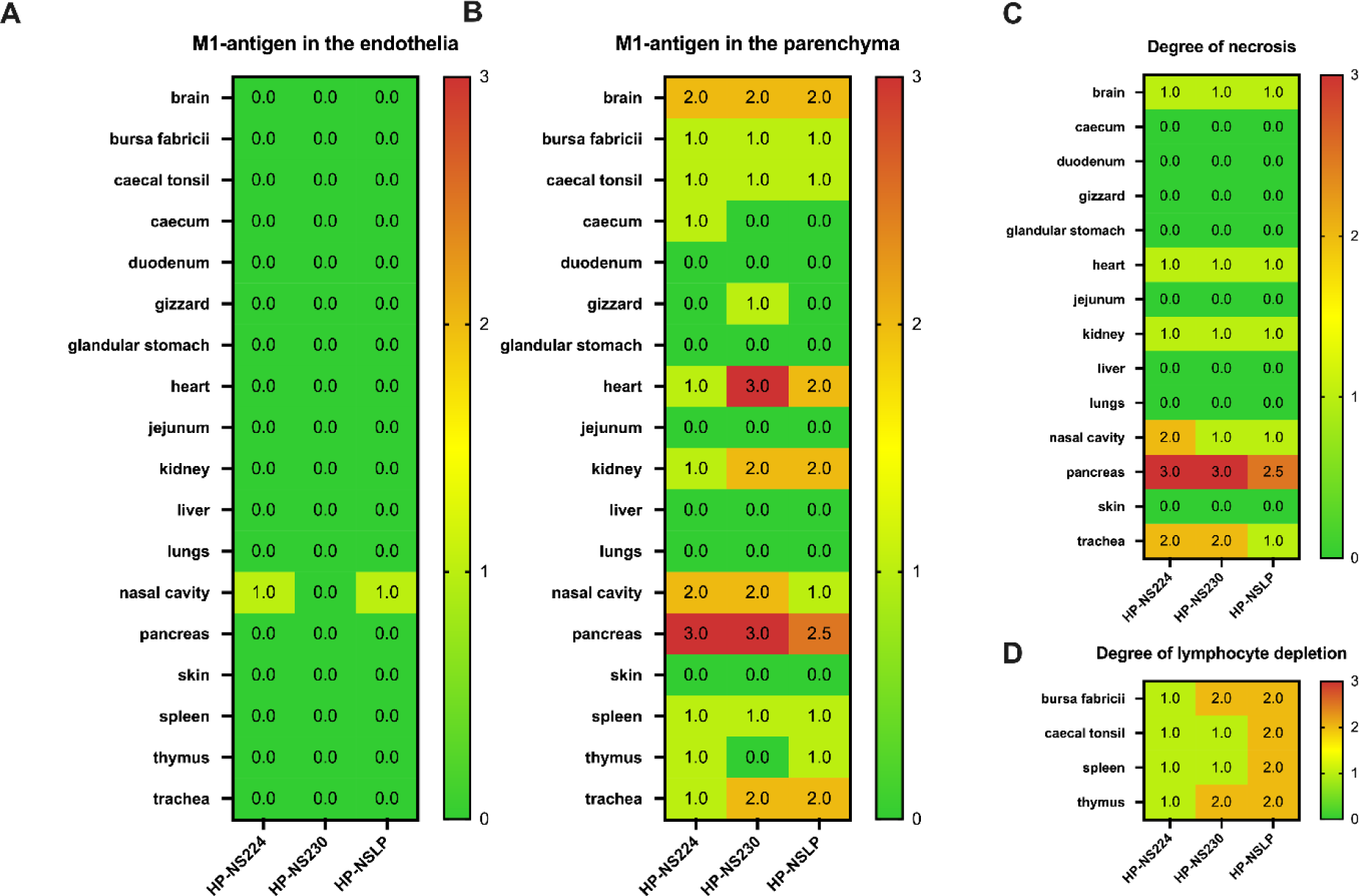
Distribution and pathological findings in H7N1-inoculated turkeys. Samples were collected from inoculated turkeys 4 dpi. M1 antigen distribution was assessed semiquantitatively in the endothelial cells (A) and parenchyma (B) of the indicated organs. The degree of necrosis (C) and lymphocyte depletion (D) was also assessed microscopically. The color of the heatmap corresponds to the median of the semiquantitative scores: 0 = no, 1 = mild, 2 = moderate, 3 = severe lesion for necrosis and depletion, 0 = no, 1 = focal to oligofocal, 2 = multifocal, 3 = coalescing to diffuse antigen for parenchyma and 0 = no antigen, 1 = antigen in single blood vessels, 2 = antigen in multiple blood vessels, 3 = diffuse immunoreactivity for endothelial cells.

**Fig. 5:**
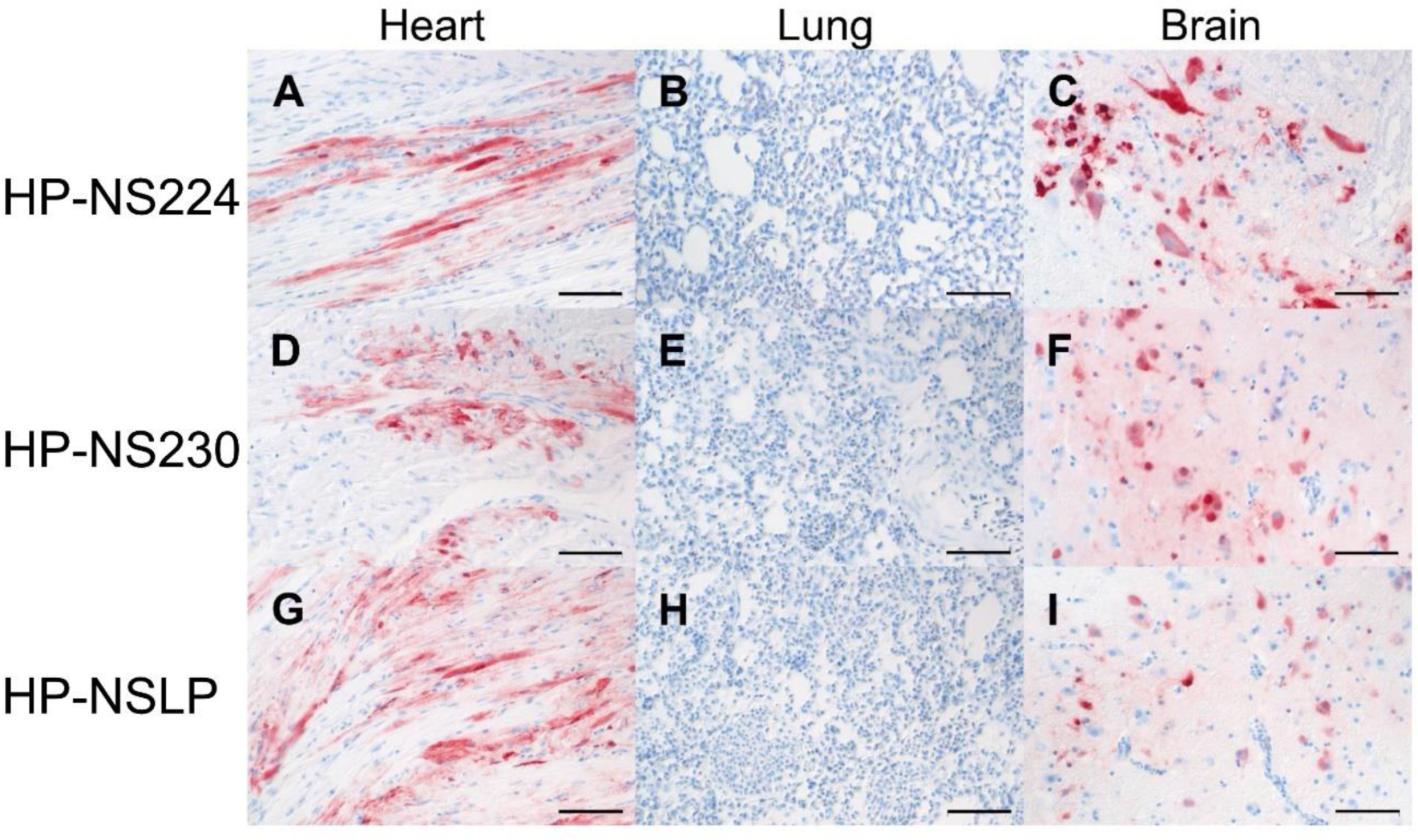
Detection of H7N1 matrix 1 protein in selected tissues of inoculated turkeys. The maximum distribution of influenza A virus M1 protein 4dpi in heart, lung and brain of turkeys inoculated with HP-NS224 (A, B, C), HP-NS230 (D, E, F) and HP-NSLP (G, H, I). Immunohistochemistry was done with the avidin-biotin-peroxidase-complex method, a primary monoclonal mouse antibody against influenza A virus matrix protein, 3-amino-9-ethylcarbazol (red-brown) as chromogen and hematoxylin (blue) as counterstain; Nomarski contrast; Bars = 50 µm.

### Co-localization of NS1 and M1 in the brain of H7N1 infected turkeys

To date, NS1 as a non-structural protein is not considered to be part of the infectious virion and no data are available on the expression or distribution of NS1 in infected birds. As a proof of principle, we sought to identify NS1 in the brain of HP-NS224 infected turkeys at 4 dpi using immunofluorescence (Fig. 6A-D). Interestingly, multiple foci of M1 and NS1 antigen expression were detected in the brain. Higher magnification showed simultaneous M1 and NS1 expression in neurons, which were clearly identifiable by their characteristic morphology (Fig. 6B). NS1 expression was equally strong in the cytoplasm and nucleus (Fig. 6C), whereas M1 expression tended to be stronger in the nucleus (Fig. 6D). The NS1 signal was stronger and better discriminated from the autofluorescent background than the M1 signal in the immunofluorescence experiment. Simultaneously processed heart, spleen and lung tissues showed overwhelming autofluorescence due to high amounts of erythrocytes, preventing reliable analysis of these tissues by immunofluorescence (data not shown). These results indicate that, similar to the M1 antigen, NS1 expression is stable in the brain of H7N1 inoculated turkeys 4 dpi and can be used for in vivo experiments (e.g. to study interaction with host factors in vivo).

**Fig. 6:**
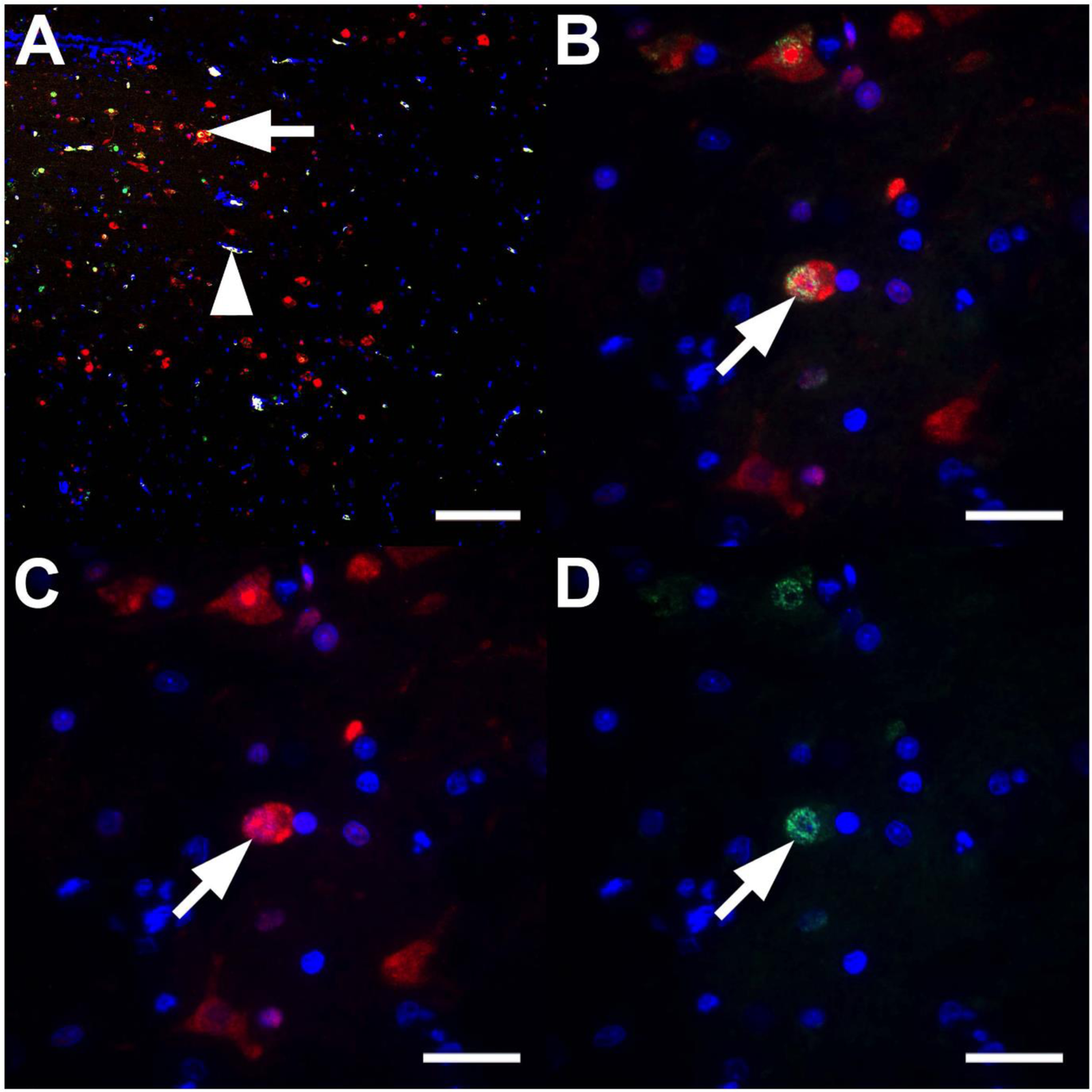
Co-localization of NS1 and M1 antigens in the brain tissues of H7N1 inoculated turkeys. Immunofluorescence was used to proof simultaneous expression of influenza A virus non-structural antigen 1 (NS1) and matrix-1 protein (M1). The brain of a HP-NS224-infected turkey 4 dpi was immunolabelled for NS1 in red (C), M1 in green (D), both with nuclei in blue. In the merged images (A-B) colocalization of NS1 and M1 is shown in yellow and nuclei in blue. (A) The merged low magnification image of the cerebrum displays a periventricular focus (ependymal nuclei lined up like a ribbon in the upper left corner) of NS1 and M1 expression, often within polygonal cells, resembling neurons, visible as yellow double-labelling (arrow) in the merged image. The arrowhead points to a vessel exhibiting intense, white, false positive autofluorescence. (B-C) A higher magnification of a group of characteristic neurons displays an equal intensity of NS1 in the cytoplasm and nucleus, as compared to a more nuclear expression of M1 (A-D) Immunofluorescence for NS1 with AlexaFlour 488-coupled secondary antibody, M1 with Alexa-456-coupled secondary antibody and nuclei with 4’,6-diamidino-2-phenylindole. (A) Scale bar = 100 µm; (B-D) scale bars = 20 µm.

## Discussion

Turkeys and chickens are Galliformes and are highly susceptible to AIV infection ^15^. Because of their higher susceptibility to AIV infection than chickens, particularly to H7 viruses ^16, 17^, turkeys play an important role in the adaptation of AIVs of wild bird origin to poultry ^18, 19^. In contrast to ducks, chickens and turkeys usually die after HPAIV infection. Therefore, it has been suggested that virulence determinants and virus-host interactions in AIV-infected turkeys are likely to be similar to those in chickens. However, we have recently shown that virulence determinants located in the HA of two different H7 viruses are different in chickens and turkeys ^20, 21^. No data are available on the role of NS1 in AIV fitness in turkeys. A few studies have shown that mutations in the NS1 of H7Nx, particularly in ED or CTE, affect virus fitness in chicken embryos^11^, chickens ^13, 22–26^ and ducks ^13, 27^.

In the current in vitro experiments, we found no significant differences in multiple-cycle virus replication or NS1 expression in turkey cells. Soubies, et al. ^13^ found that the 6-aa truncation associated with transition of the Italian LPAIV H7N1 to HPAIV had no effect on LP virus replication in duck or chicken in vitro and in vivo. Similarly, all three NS1s were equally effective in blocking type I IFN mRNA induction in chicken and human cell lines, which is consistent with the findings of Soubies, et al. ^13^ who described comparable levels of type I IFN mRNA induced by similar H7N1 LP carrying NS230-aa or NS224-aa in chicken and duck cells. Conversely, Keiner, et al. ^11^ found that prolongation of the CTE of HPAIV H7N1 NS1 reduced the efficiency of blocking IFN mRNA induction using a different cell culture and methodology to that used in the current study. In our previous chicken experiment using the same three viruses, we found that NS1 elongation reduced virus replication in tissues of infected chickens ^10^. Conversely, in turkeys we did not observe significant variation in the distribution of the H7N1 in different organs, further ruling out a role for NS1 as a virulence determinant in turkeys ^21^, in contrast to chickens or chicken embryos ^11, 21^.

Currently, it is widely accepted that NS1 is a non-structural protein that is only expressed in cells after infection and is not part of the virion. Anti-NS1 antibodies have been found in the sera of birds experimentally infected with different AIV subtypes, suggesting that NS1 antibodies could be used for diagnostic purposes ^28–30^. However, no studies, so far, have reported the detection of NS1 antigen specifically in the brain of turkeys. Our pilot study showed that NS1 antigen can be detected in the turkey brain, similar to and overlapping with M1, indicating that the virus is replicating in the neurons. This will be useful in subsequent experiments to further improve our understanding of the potential role of NS1 *in vivo*, for example, in neurovirulence or blocking the immune response in the turkey brain.

In conclusion, we found that turkeys can be infected with AIV carrying different NS1 lengths and mutations without significant effects on multiple-cycle replication, and NS1 expression in vitro, and without affecting the high virulence, efficient transmission and tissue distribution of HPAIV H7N1 in turkeys. These results are in contrast to our previous findings in chickens, where NS1 significantly affected virus replication in vitro and in vivo. Our results further suggest that the pathogenesis and genetic markers for adaptation of AIV are different in chickens and turkeys, although they are very closely related galliform species.

## Material and methods

### Sequence analysis

NS1 sequences of H7N1 isolated from turkeys and chickens in Italy in 1999 and 2000 were retrieved from GISAID on 16-01-2024. Sequences with double entries and identical amino acid sequences were further edited. Alignment was done using Geneious Prime® Software (Version 2021.0.1) and the MAFFT package.

### Cells and recombinant viruses

Primary turkey embryonic kidney cells (TEK) were prepared from the kidneys of 21-day old turkey embryos ^21, 31^. Human embryonic kidney 293T (HEK-293T), Madin-Darby canine kidney type II (MDCKII) and chicken fibroblast (DF1) cell lines were obtained from the Cell Bank of Friedrich-Loeffler-Institut (FLI). For this study, previously generated recombinant HPAIV H7N1 were used ^10^: A/chicken/Italy/445/1999 virus carrying the HP NS1 (HP-NS224), the LP NS1 form A/chicken/Italy/473/1999 virus (LP-NSLP) and the elongated HP NS1 (HP-NS230) ^10^. Virus stocks were prepared in embryonated chicken eggs (ECE) obtained from specific-pathogen-free chickens (VALO BioMedia GmbH, Germany). Sequence of the whole genome of the three viruses was determined as previously done ^10^.

### Plasmids

pCAGGS plasmids carrying the NS inserts of HP-NS224, HP-NS230, and HP-NSLP were generated in this study. The NS segment was amplified using the Omniscript RT Kit (Qiagen, Germany) and Phusion® High-Fidelity DNA Polymerase (New England Biolabs, USA). PCR products were purified on a 1% agarose gel and extracted using the GeneJET G el Extraction Kit (Thermo Fisher Scientific, Germany). The competent E. coli XL1-blue strain was transformed and plasmids were isolated from bacterial cultures using the Plasmid Midi Kit (Qiagen, Germany). Sequencing of the three NS1 plasmids was performed by Eurofins (Germany).

### Luciferase reporter assay

To determine the efficiency of NS1 in inhibiting the IFN-I pathway, we conducted luciferase reporter assay as previously described [<u>5</u>]. Briefly, HEK293T and DF1 cells in 6 well plates were transfected with a plasmid DNA mixture containing 0.5 μg of a Firefly Luciferase (FFL) expressing reporter plasmids (i.e. firefly luciferase reporter plasmids with human or chicken IFN-I promotors:pIFN-ß-Pro-FFL, pIFNa-Pro-FFL), 0.005 μg of pCMV-RL (normalization), 0.2 μg human or chicken pIRF7 or 0.5 μg human or chicken pMDA5-delta or human pTrif as a trigger expression plasmid and 0.5 μg of pCAGGS plasmid with one of the NS1 coding sequences or an empty vector as a control. Lipofectamine™ 2000 (Thermo Fisher) was used for the transfection according to manufacturer recommendations. At 24 h post-transfection, cell lysates were harvested and luciferase activity was measured using the Dual-Luciferase® Reporter Assay System (Promega, USA) according to the manufacturer’s instructions. Firefly and Renilla activity were measured using a TriStar² S LB 942 Modular Multimode Microplate Reader (Berthold, Germany). The assay was done in three independent experiments and results are expressed as normalized means and standard deviations.

### Western blot

TEK cells, 80% confluent, were infected with viruses (MOI 0.1) for 1h, washed twice with phosphate-buffered saline (PBS) and overlaid with Ham’s F12/IMDM supplemented with 0.2% bovine serum albumin (BSA, MP Biomedicals, USA). Cells were harvested at 24 hours post infection (hpi). Samples were centrifuged at 10,000 rpm for 5 minutes, washed with 0.5 ml PBS, centrifuged at 10,000 rpm for 10 minutes, resuspended in 50 µl of PBS and 50 µl of Laemmli sample buffer 2x, for SDS-PAGE (Serva, Germany), boiled at 99°C for 10 minutes and stored at -20°C until further use. Samples were run on 12% SDS-PAGE gel at 200V for 47 minutes, blotted with BioRad Turbo Blotter, blocked with 5% milk overnight. They were stained with rabbit anti-NS1 polyclonal antibody (kindly provided by Daniel Marc, INRAE, Nouzilly, France), rabbit anti-NP polyclonal antibody and GAPDH as a cell normalization control (Abcam, United Kingdom). Blots were developed using the Biorad VersaDoc system and quantified using Image J software.

### Replication kinetics

Recombinant viruses were assayed for growth rate by infecting TEK cells at MOI of 0.001 for 1h. The cells were washed twice with PBS and overlaid with Ham’s F12/IMDM medium supplemented with BSA. Plates containing cells were incubated at 37°C in 5% CO2 for 1, 8, 24, 48 and 72 hours. Prior to titration by plaque assay, harvested cells were stored at -80°C.

### Plaque assay

The plaque assay was used for virus titration and virus titers were expressed as plaque-forming units per mL (PFU/mL). Ten-fold serial dilutions of each virus were incubated on MDCKII cells in 6-well plates for one hour at 37°C and 5% CO2, washed twice with PBS, 0.9% Bacto Agar/MEM mixture supplemented with 0.2% BSA (MP Biomedicals, USA) and incubated for 72 hours at 37°C and 5% CO2. After incubation, the plates were fixed with 0.1% crystal violet in 10% formaldehyde solution. Viral titers were determined by counting the number of plaques under a microscope.

### Turkey experiment. Ethical approval

The animal experiment was performed in the Biosafety Level 3 (BSL3) facility of the FLI in accordance with the German Animal Welfare Act after approval by the authorized Ethics Committee of the State Office for Agriculture, Food Safety and Fisheries of Mecklenburg-Western Pomerania (LALLF M-V) under registration number 7221.3-1.1-051-12. A total of 45 six-week-old turkeys were purchased from a farm in Mecklenburg-Western Pomerania. Each recombinant virus was used to oculonasally infect 10 birds per group at 10^4.5^ PFU/bird. After 24 h, 5 naive birds were added to each group. All birds were observed daily for clinical signs (depression, signs of respiratory distress, diarrhea, cyanosis of the comb, wattles or shanks, facial edema and neurological signs) over a 10-day observation period. Clinical scoring followed the standard protocol ^33^ where healthy birds were scored (0), sick birds with one of the clinical signs were scored (1), severely sick birds with two or more signs were scored (2) and dead birds were scored (3). Moribund birds unable to eat or drink were humanely euthanized by inhalation of isoflurane (CP-Pharma, Germany), exsanguinated and scored as dead on the next observation day. The pathogenicity index (PI) was calculated as the sum of the daily arithmetic means of all infected birds divided by 10 (the number of observation days), with a final range from 0 (avirulent) to 3 (highly virulent).

### Virus shedding

Oropharyngeal and cloacal swabs were taken from all birds at 2 and 4 dpi. Swabs were stored at -80°C in 1.5 ml Dulbecco’s Modified Eagle Medium (DMEM) containing 1.05 mg enrofloxacin (Baytril, Bayer AG, Germany), 0.525 mg lincomycin (Mediserv Nord, Germany), 0.105 mg gentamycin (Genta, CP-Pharma, Germany) in sterile safe-lock Eppendorf tubes, previously mixed by vortexing for 30 seconds. The amount of infectious virus in plaque-forming units was determined by titration of the swab samples using the plaque assay described above.

### Histopathology and immunohistochemistry

For histopathology and immunohistochemistry, organ samples were fixed in 4% neutral buffered formaldehyde for >7 days, processed, embedded in paraffin and sectioned at 2-4 μm. For histopathology, slides were stained with haematoxylin and eosin. Immunohistochemistry was performed using the avidin-biotin-peroxidase complex method (Vectastain PK 6100; Vector Laboratories, Burlingame, CA, USA) with citric buffer (pH 6.0), a primary mouse monoclonal antibody against influenza A virus matrix 1 protein (M1, ATCC clone M1Hb-64, 1:100), a secondary biotinylated goat anti-mouse IgG (BA-9200, Vector Laboratories, Newark, USA, 1:200), 3-amino-9-ethylcarbazol as chromogen, and hematoxylin counterstain as described [<u>13</u>, <u>18</u>]. Mouse IgG (NBP1-97019-5mg, Novus Biologicals USA, CO, USA) was used as an isotype control instead of the primary antibody and validated archival tissues were used as a positive control.

Scoring of histopathological lesions and viral antigen distribution was performed as previously described ^36^. Briefly, necrosis or necrotizing inflammation and lymphoid depletion were each scored as follows 0 = no, 1 = mild, 2 = moderate, 3 = severe lesion, and viral antigen distribution was scored as follows: 0 = no, 1 = focal to oligofocal, 2 = multifocal, 3 = coalescing to diffuse antigen for parenchymal cells and 0 = no antigen, 1 = antigen in single blood vessels, 2 = antigen in multiple blood vessels, 3 = diffuse immunoreactivity for endothelial cells.

### Immunofluorescence double-staining

An immunofluorescence double-staining experiment was performed for proof-of-principle of influenza A virus matrix 1 protein (M1) and non-structural protein 1 (NS1) co-expression on the brain of a HP-NS224-infected turkey (same animal as shown in Fig. 5 A-C), as well as a H4/H5-infected chicken, and a non-infected chicken from previous studies as positive and negative controls, respectively. Briefly, 2-4 µm slides of the formalin-fixed paraffin-embedded specimen were deparaffinized, and sequentially incubated in 0.5% H2O2 in methanol for 30 minutes for inhibition of endogenous peroxidase, in citric acid buffer (pH 6,0) at 96°C for 25 minutes for unmasking of antigens, in 0,1 % Triton X-100 in Tris buffered saline (TBS) for 15 minutes for permeabilization, and SuperBlock^TM^ blocking buffer (Thermo Fisher Scientific, USA) for 30 minutes for blocking excess binding sites. Afterwards, the slides were incubated with either single or mixed primary antibodies against M1 (monoclonal mouse anti-M1 antibody (ATCC, clone: M1Hb-64, 1:10) and NS1 (polyclonal anti-rabbit, 1:1000) at 4°C overnight, followed by incubation with AlexaFluor 488-conjugated donkey anti-mouse IgG (1:500, Dianova) and AlexaFluor 568-conjugated goat anti-rabbit IgG (1:500, Abcam) secondary antibodies at room temperature for 1 h. After further washing steps, sections were counterstained using 4’,6-diamidino-2-phenylindole (DAPI, 1:300, Invitrogen) and mounted with glycerol-gelatin aqueous slide mounting medium (Sigma Aldrich). The labelled sections were analyzed using a motorized Axioplan 2 Imaging fluorescence microscope (Carl Zeiss Microscopy Deutschland GmbH, Oberkochen) equipped with 25x/0.8 Plan-Neofluar water-immersion, 40x/1,2 Apochromat water-immersion, and 63x/1,2 Apochromat water-immersion objectives, an HBO 50 mercury-vapor short-arc lamp, AHF F31-000 (excitation 350/50, emission 460/50, for DAPI), AHF F41-054HQ (excitation 480/30, emission 527/30 HQ, for AF488), and AHF F41-007HQ (excitation 545/30, emission 610/75 HQ, for AF568) filter sets, and a monochrome 12 megapixel Axiocam 712 mono R2 CMOS camera. Images were acquired using a semi-automated multi exposure protocol within the ZEN 3.8 software.

### Statistic

Statistical analysis was performed using Graph Pad Prism, Version 10.1.1. One-way ANOVA was used for analysis of viral titers, expression levels of NS1 and IFN-induction inhibition. Kruskal-Wallis tests and Mann-Whitney-Wilcoxon tests with Benjamini-Hochberg correction were used for clinical scoring. Survival analysis was done by log-rank (Mantel-Cox) test.

## Acknowledgements

The authors thank Frank Klipp, Doreen Fiedler, Charlotte Schröder, Diana Palme, Luise Hohensee, David Scheibner and colleagues at the department of experimental animal facilities and biorisk management, for their valuable assistance with the animal experiments. Timm C. Harder, FLI and Ilaria Capua, Insituto Zooprofilattico Sperimentale delle Venezie, Padova, Italy, for providing the viruses. Dajana Helke, Kristin Trippler, Elfi Quente and Hilke Gräfe for technical assistance. We thank Daniel Marc for the anti-NS1 antibodies and Stefan Finke for the pCAGGS expression vector.

## Data availability

Data will be made available on request

## Author contributions

Conceptualization: EMA, Data curation: All authors, Formal analysis: All authors Funding acquisition: EMA, RU, CL, Investigation: All authors, Methodology: All authors, Project administration: EMA, Resources; EMA, RU, CL, Software; All authors, Supervision: EMA, RU, Validation; All authors, Visualization; All authors, Roles/Writing - original draft: MK, EMA and Writing - review & editing: All authors.

